# Spectrin is a mechanoresponsive protein shaping the architecture of intercellular invasion

**DOI:** 10.1101/154831

**Authors:** Rui Duan, Ji Hoon Kim, Khurts Shilagardi, Eric Schiffhauer, Sungmin Son, Donghoon Lee, Shuo Li, Graham Thomas, Tianzhi Luo, Daniel A. Fletcher, Douglas N. Robinson, Elizabeth H. Chen

## Abstract

Spectrin is a membrane skeletal protein best known for its structural role in maintaining cell shape and protecting cells from mechanical damage^1-3^. Here, we report that spectrin dynamically accumulates and dissolves at the fusogenic synapse, where an attacking fusion partner mechanically invades its receiving partner with actin-propelled protrusions to promote cell-cell fusion^4-7^. Using genetics, cell biology, biophysics and mathematical modeling, we demonstrate that unlike myosin II that responds to dilation deformation, spectrin exhibits a mechanosensitive accumulation in response to shear deformation, which is highly elevated at the fusogenic synapse. The accumulated spectrin forms an uneven network, which functions as a “sieve” to constrict the invasive fingerlike protrusions, thus putting the fusogenic synapse under high mechanical tension to promote cell membrane fusion. Taken together, our study has revealed a previously unrecognized function of spectrin as a dynamic mechanoresponsive protein that shapes the architecture of intercellular invasion. These findings have general implications for understanding spectrin function in other dynamic cellular processes beyond cell-cell fusion.

---

The mechanical properties of cells are dynamically controlled in many cellular processes, such as cell division, fusion, migration, invasion, and shape change. Spectrin is best known as a membrane skeletal protein critical for maintaining cell shape and providing mechanical support for plasma membrane^1-3^. In erythrocytes and neurons, spectrins, together with actin, ankyrin and associated proteins, form a static polygonal lattice structure^8-10^ or an ordered periodic longitudinal array^11^ underneath the plasma membrane to protect cells from mechanical damage^12^. Such a mechanoprotective function of spectrin is made possible by holding the spectrin network under constitutive tension^13^. However, in many cellular processes, mechanical tension is generated upon transient cell-cell interactions. How spectrins, which are expressed in most eukaryotic cells, respond to transient and localized mechanical stimuli in dynamic cellular processes remains largely unknown.

Cell-cell fusion is a dynamic process that occurs in fertilization, immune response, bone resorption, placenta formation, and skeletal muscle development and regeneration^14,15^. It has been demonstrated in a variety of cell fusion events from *Drosophila* to mammals that cell fusion is an asymmetric process^4-7,16^. At the site of fusion, known as the fusogenic synapse, an attacking fusion partner invades its receiving fusion partner with actin-propelled membrane protrusions^4-7,16^, while the receiving fusion partner mounts a myosin II (MyoII)-mediated mechanosensory response^6^. The invading and resisting forces from the two fusion partners, in turn, push the two cell membranes to close proximity and put the fusogenic synapse under high mechanical tension to promote fusogen engagement and cell membrane merger^5,6^. Although previous studies have shown that multiple long (1-2 μm) and thin (~200 nm in diameter) invasive protrusions from the attacking fusion partner are required for cell-cell fusion^4,5,17,18^, it is unclear how these protrusions are spatially constricted and shaped in order to provide high mechanical tension at the fusogenic synapse.

In a genetic screen for new components required for cell-cell fusion, we have uncovered an essential function for spectrin in *Drosophila* myoblast fusion. Spectrins are composed of α- and β-subunits, which laterally associate to form antiparallel heterodimers and then assemble head-to-head to form flexible, rod-like heterotetramers with actin-binding domains at the two ends^1-3^. The *Drosophila* genome encodes one α-, β- and β_heavy(H)_-subunit, the latter of which (also known as *karst*) is homologous to βV-spectrin in vertebrates^19^. Zygotic mutants of α- or β_H_-spectrin exhibited minor myoblast fusion defects (Fig. 1a-h, i) likely due to maternal contribution, whereas α/β*_H_-spectrin* double mutant showed a severe fusion defect (Fig. 1e, i). The functional specificity of α/β_H_-spectrin in myoblast fusion was demonstrated by a genetic rescue experiment, in which full-length β_H_-spectrin expressed in all muscle cells rescued the myoblast fusion defect in β*_H_-spectrin* mutant embryos (Fig. 1f, i), and by an overexpression experiment, in which a dominant negative form of β_H_-spectrin (mini-β_H_-spectrin, deleting 15 of the 29 spectrin repeats)^20^ expressed in muscle cells enhanced the fusion defect of β*_H_-spectrin* mutant embryos (Fig. 1g, i). Furthermore, expressing β_H_-spectrin specifically in the receiving fusion partners, the muscle founder cells, rescued the fusion defect (Fig. 1h, i), demonstrating that α/β_H_-spectrin functions specifically in the receiving cells.

**Figure 1.**
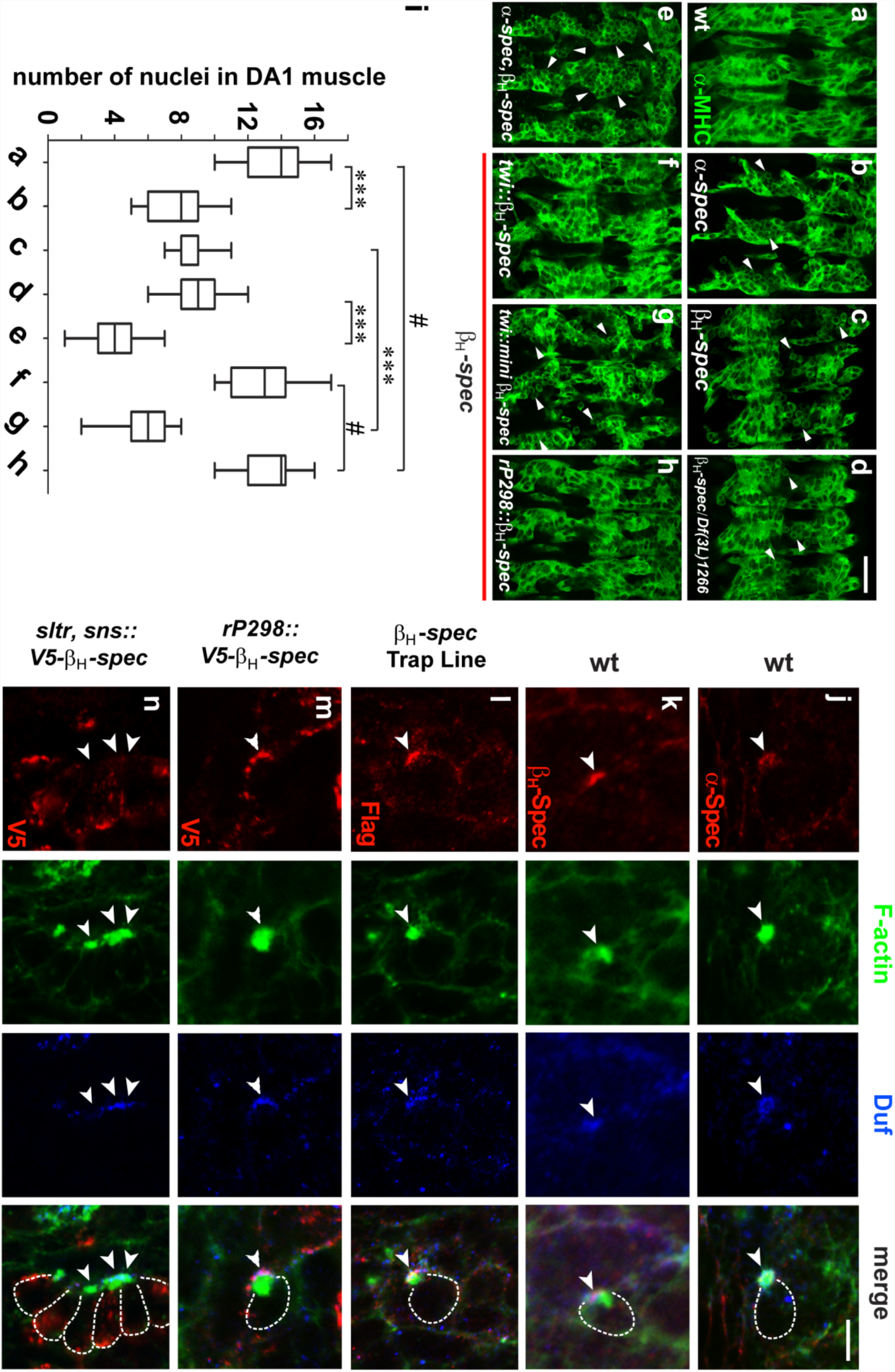
Spectrin is required for myoblast fusion and enriched at the fusogenic synapse in the receiving cell. (a-h) Myoblast fusion phenotype in *α/β_H_-spectrin* mutant embryos. Somatic muscle cells in stage 15 embryos were labeled with anti-muscle myosin heavy chain (MHC) antibody. Ventral lateral muscles of three hemisegments are shown in each panel. Anterior is to the left and posterior to the right. (a) Wild-type (wt). (b-d) Minor fusion defect in *α-spectrin* (*α-spec*) (b), *β_H_-spectrin* (*β_H_-spec*) (c), and transheterozygous *β_H_-spec/Df(3L)1266* (d) mutant embryos. (e) Severe fusion defect in *α/β_H_-spectrin* double mutant embryo. (f-h) Effect of *β_H_-spec* transgenes expression in *β_H_-spec* mutant embryos. Expression of full-length *β_H_-spec* in all muscle cells with a *twi-GAL4* driver (f) or in founder cells with an *rP298-GAL4* driver (h) rescued the fusion defect. Expression of *mini-β_H_-spec*, a dominant negative form of β_H_-Spec, in all muscle cells enhanced the fusion defect (g). Unfused cells are indicted by arrowheads. Scale bar, 20 μm. (i) Quantification of the fusion index. The number of nuclei in the dorsal acute muscle 1 (DA1) were counted for each genotype in (a-h). Error bars indicate standard deviation. Statistical significance was determined by the Student’s t test. ***, p < 0.001; #, not significant. (j-n) Localization of α/β_H_-spectrin at the fusogenic synapse. Stage 13-14 embryos were triple labeled with phalloidin (F-actin; green), anti-Duf (founder cell adhesion molecule; blue), as well as anti-α-Spec (j), anti-β_H_-Spec (k), anti-Flag (β_H_-spec-Flag trap; l), anti-V5 (V5-β_H_-Spec expressed in founder cell; m) and anti-V5 (V5-β_H_-Spec expressed in FCM with an *sns-GAL4* driver; n). Note the accumulation of α-Spec (j) and β_H_-Spec (k and l) at the fusogenic synapse and their colocalization with Duf. The white outlines in the merged panels delineate FCMs which attacked founder cells at the fusogenic synapse. Scale bar, 5 μm. (m-n) Founder cell-specific accumulation of β_H_-Spec at the fusogenic synapse. V5-tagged β_H_-Spec was specifically expressed in founder cells (m) or in FCMs (n) and visualized by anti-V5 staining. Expression of β_H_-Spec in FCMs was performed in a fusion-defective *sltr* mutant to prevent diffusion of ectopically expressed β_H_-Spec proteins from FCMs to founder cells following myoblast fusion events. Note that β_H_-Spec was accumulated in founder cells (m), not in FCMs (n), at the fusogenic synapse.

To determine the subcellular localization of α/β_H_-spectrin, we performed antibody-labeling experiments using anti-α- and β_H_-spectrin antibodies in wild-type embryos (Fig. 1j, k), and an anti-Flag antibody in a protein trap line, *kst^MI03134-GFSTF.1^*, which carries 3xFlag-tagged β_H_-spectrin (Fig. 1l). Both α- and β_H_-spectrin are highly enriched at the fusogenic synapse marked by an F-actin-enriched focus, whereas no enrichment was observed for β-spectrin, which was expressed at a high level in epithelial cells (Extended Data Fig. 1a). The β_H_-spectrin enrichment at the fusogenic synapse largely colocalized with that of Dumbfounded (Duf), an Ig domain-containing founder cell-specific adhesion molecule^21^ (Fig. 1j-l). Moreover, expressing a V5-tagged β_H_-spectrin specifically in founder cells led to the accumulation of β_H_-spectrin-V5 at the fusogenic synapse (Fig. 1m). In contrast, expressing β_H_-spectrin-V5 specifically in the attacking fusion partners, the fusion competent myoblasts (FCMs), did not result in its accumulation at the fusogenic synapse (Fig. 1n). These results further support the functional requirement for α/β_H_-spectrin in founder cells.

To investigate whether β_H_-spectrin forms a stable membrane skeletal network at the fusogenic synapse, we performed live imaging experiments in *Drosophila* embryos. Surprisingly, instead of forming a stable network, β_H_-spectrin exhibited dynamic accumulation and dissolution at the fusogenic synapse accompanying the appearance and disappearance of the actin-enriched invasive structure from the FCM, known as a podosome-like structure (PLS)^4^ (Supplementary Video 1; Extended Data Fig. 2a, b). Previous studies have shown that the density of an actin focus correlates with its invasiveness, such that loosely packed actin filaments are less invasive, as in *dpak3* mutant embryos^18^. We found that the amount of β_H_-spectrin accumulation correlated with the density of the F-actin foci, as shown in both wild-type (Supplementary Video 1; Extended Data Fig. 2a, b) and *dpak3* mutant embryos (Supplementary Video 2; Extended Data Fig. 2c, d). Specifically, β_H_-spectrin accumulation in *dpak3* mutant embryos was overall weaker than that in wild-type embryos, except for a few spectrin “hot spots” associated with the occasionally aggregated small F-actin foci (Extended Data Fig. 2c, d). These findings suggest that spectrin does not form a static network during myoblast fusion; instead, it forms a transient and dynamic structure that rapidly changes its density, shape and morphology in response to changes in the invasiveness of the PLS. The dynamic behavior of β_H_-spectrin at the fusogenic synapse was confirmed by fluorescent recovery after bleaching (FRAP). When fluorescently labeled β_H_-spectrin was photobleached at the fusogenic synapse as its accumulation reached equilibrium, the fluorescence rapidly recovered with an average T_1/2_ of 59 ± 18 sec (n=6) and eventually reached 54 ± 9% (n=6) of the pre-bleaching level (Fig. 2a, b; Supplementary Video 3). These results suggest that more than half of β_H_-spectrin at the fusogenic synapse is in a mobile fraction and that β_H_-spectrin dynamically associates and dissociates at the fusogenic synapse, rather than being maintained in a stagnant structure.

**Figure 2.**
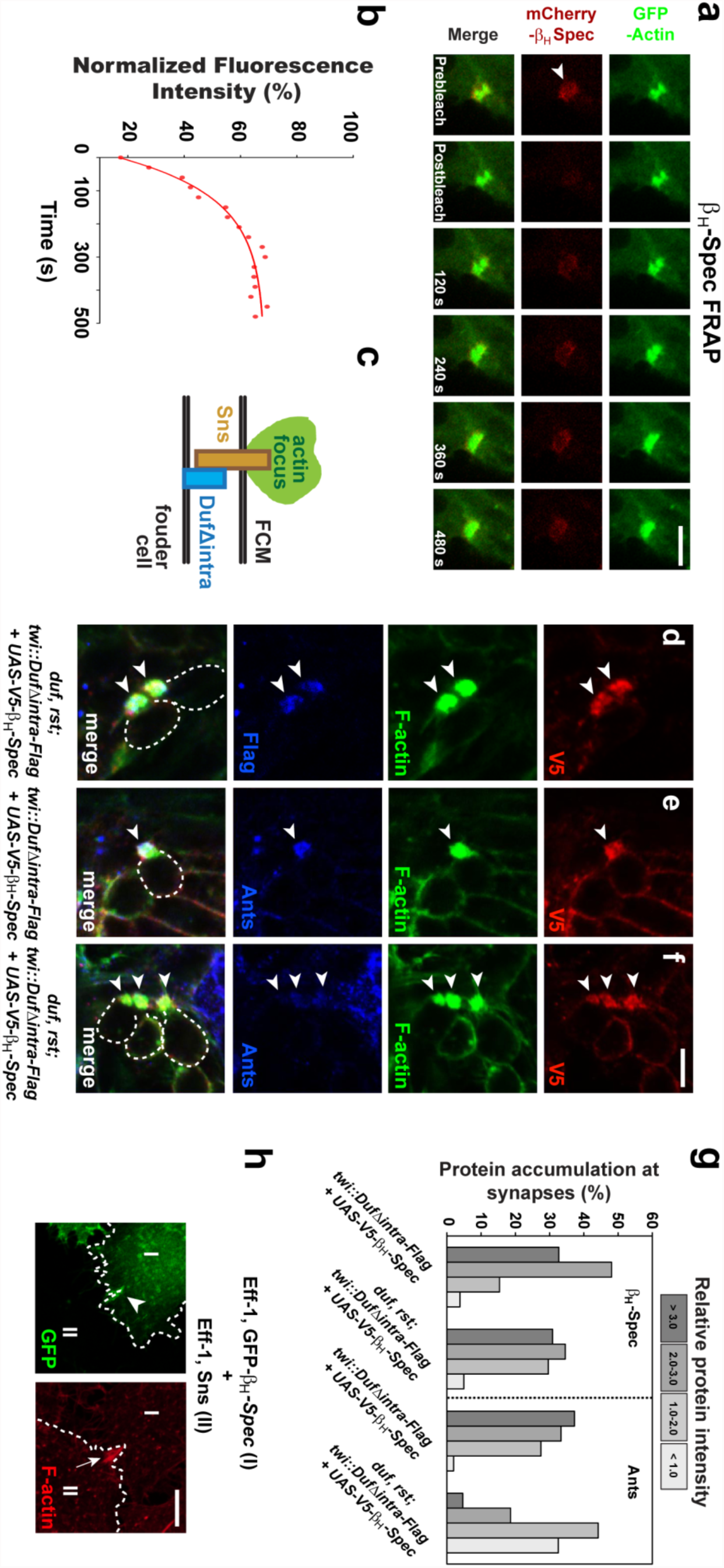
Spectrin dynamically accumulates at the fusogenic synapse in the absence of chemical signaling from cell adhesion molecules. (a-b) FRAP of β_H_-Spec at the fusogenic synapse. (a) Stills from time-lapse imaging of a stage 14 wt embryo expressing GFP-actin and mCherry-β_H_-Spec. Arrowhead indicates the photobleached region. Scale bar, 5 μm. (b) Recovery kinetics of the mCherry fluorescence after photobleaching. (c-g) β_H_-Spec accumulation at the fusogenic synapse in the absence of chemical signaling. (c) A schematic drawing of the fusogenic synapse showing truncated Duf (DufΔintra) in the founder cell interacting with Sns in FCM to induce the formation of an invasive F-actin focus. (d-f) Co-expression of V5-β_H_-Spec (red) and DufΔintra-Flag (blue) in all muscle cells of *duf,rst* double mutant (d and f) and wt (e) embryos. The FCMs are outlined in the merge panels. Note the β_H_Spec accumulation (d and f) and the lack of accumulation of Ants (anti-Ants; blue; f) at the fusogenic synapse (arrowheads) in the absence of Duf intracellular signaling. (g) Quantification of the relative intensity of β_H_-Spec and Ants enrichment at the fusogenic synapse in wt and DufΔintra-expressing *duf,rst* double mutant embryos. The fluorescent intensity at the fusogenic synapse was compared with that in the adjacent cortical area to calculate the relative protein intensity ratio (n>40 for each genotype). Scale bar, 5 μm. (h) β_H_-Spec enrichment at the fusogenic synapse in fusing S2R+ cells. GFP-β_H_-Spec expressed in a receiving cell (I; outlined) accumulated at the fusogenic synapse (arrowhead) in response to an F-actin-propelled invasive protrusion (arrow) from an attacking cell (II; outlined). Scale bar, 5 μm.

Given the correlation between spectrin accumulation and PLS invasiveness, we tested whether β_H_-spectrin accumulation at the fusogenic synapse is triggered by mechanical force generated by the invasive PLS or is recruited by the cell adhesion molecules in the founder cell, Duf and its functionally redundant paralog Roughest (Rst)^21,22^. We expressed a truncated form of Duf lacking its entire intracellular domain (DufΔintra) (Fig. 2c) in *duf,rst* double null mutant embryos and asked if β_H_-spectrin could still accumulate. Remarkably, β_H_-spectrin exhibited significant accumulation at the fusogenic synapse marked by the invasive F-actin foci and DufΔintra enrichment, albeit with a slight decrease in intensity compared with the wild-type control (Fig. 2d, f, g). In contrast, Antisocial (Ants)/Rols7, a founder cell adaptor protein that binds to the intracellular domain of Duf^23-26^, did not accumulate at the fusogenic synapse (Fig. 2, compare e and f; Fig. 2g). Thus, β_H_-spectrin accumulation in the founder cell can be triggered by invasive force from the PLS, independent of the chemical signaling from cell adhesion molecules. Consistent with these results, when cultured *Drosophila* S2R+ cells were induced to fuse by expressing the FCM-specific adhesion molecule Sticks and stones (Sns) and the *C. elegans* fusogen Eff-1^5^, β_H_-spectrin also accumulated to the fusogenic synapse in response to the invasive protrusions, despite the lack of endogenous Duf and Rst in these cells^5^ (Fig. 2h).

To test directly whether β_H_-spectrin exhibits mechanosensitive accumulation, we performed micropipette aspiration (MPA) assays, in which a pulling force is applied to the *Drosophila* S2 cell cortex by a micropipette with a diameter of ~5 μm. Previously, we have shown that myosin II (MyoII) is a mechanosensory protein that accumulates to the tip of an aspirated S2 cell where the mechanical stress is high^6^. Interestingly, GFP-β_H_-spectrin also showed mechanosensitive accumulation, although unlike MyoII, GFP-β_H_-spectrin accumulated to the base area of the aspirated portion of the cell (Fig. 3a, d). This effect was not due to an increase in membranous materials in this area, since an RFP-tagged PH domain^27^, which interacts with phospholipids on the plasma membrane, did not accumulate at the base (Fig. 3b, d). Nor did this effect depend on the adaptor protein Ants that did not show mechanosensitive accumulation (Fig. 3b, d). In addition, no accumulation was observed for GFP-β_H_-spectrin-ΔC (Extended Data Fig. 3a), which deleted a C-terminal fragment containing the tetramerization domain^28^, or GFP-β_H_-spectrin-ΔN (Extended Data Fig. 3a), which deleted an N-terminal fragment containing the actin-binding domain^28^ (Fig. 3b, d; Extended Data Fig. 3b), suggesting that the mechanosensitive accumulation of β_H_-spectrin requires its tetramerization and actin-binding activities. Notably, β_H_-spectrin enrichment at the base is time- and force-dependent, which increased linearly over time until reaching its peak level at 80-90 sec after the onset of aspiration (Extended Data Fig. 4a) and increased proportionally to applied pressure (Extended Data Fig. 4b). These results suggest that α/β_H_ spectrin binding to the actin network depends on the number of binding sites at a given time rather than additional cooperative activity of the previously bound tetramers, and that the mechanical force applied to the cortical actin network leads to an increase in the binding sites for the α/β_H_ spectrin heterotetramers.

**Figure 3.**
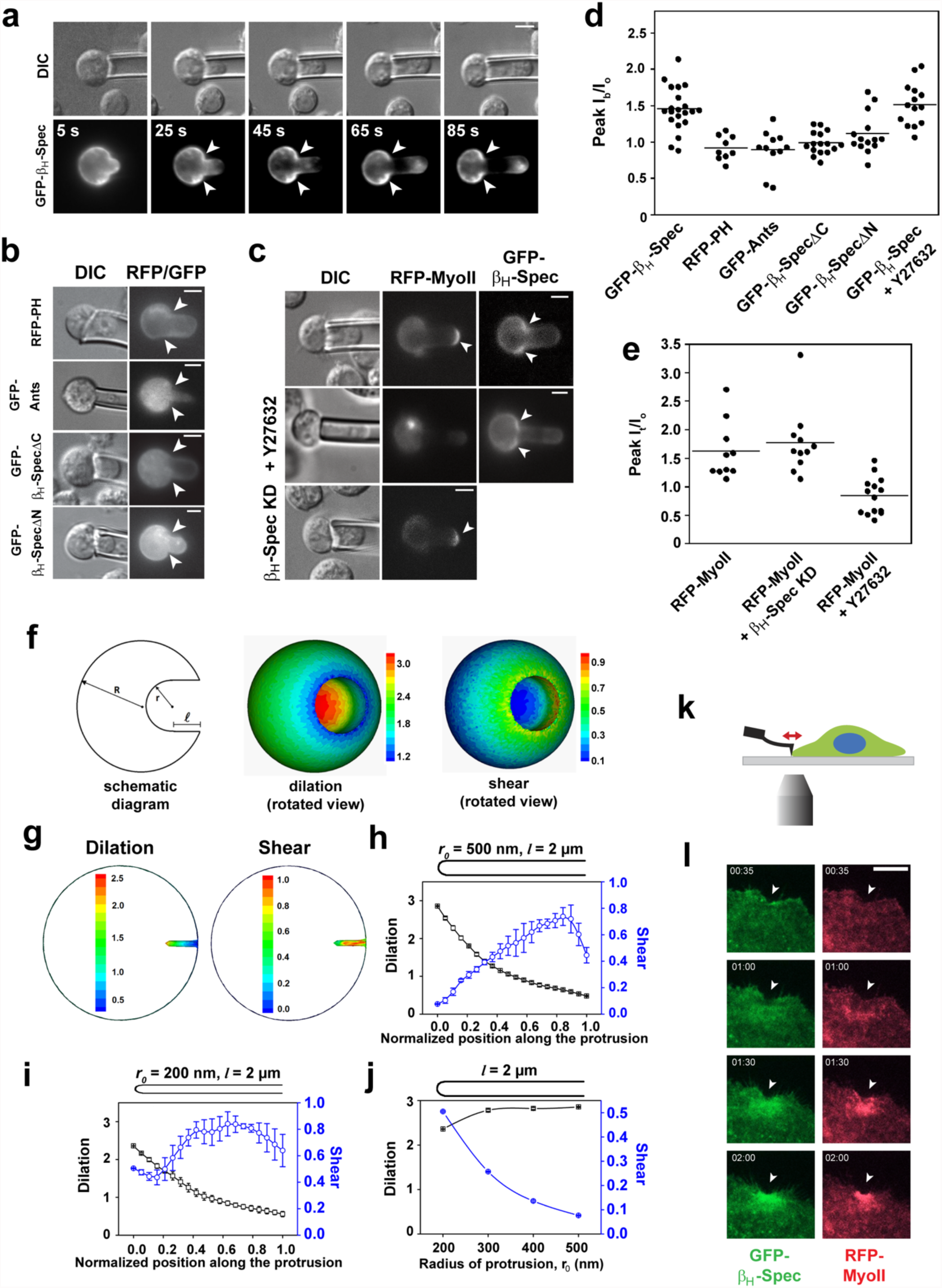
Spectrin exhibits mechanosensitive accumulation to areas of shear deformation. (a-e) Mechanosensitive accumulation of β_H_-Spec revealed by MPA. (a) Stills of time-lapse imaging of an aspirated S2 cell expressing GFP-β_H_-Spec. Differential interference contrast (DIC) and fluorescent images are shown in the top and bottom panels, respectively. Note the enrichment of β_H_-Spec to the base area (arrowheads) of the aspirated portion of the cell. (b) No mechanosensitive accumulation of a RFP-PH domain (membrane marker), GFP-Ants, GFP-β_H_-SpecΔC and GFP-β_H_-SpecΔN. (c) Spatially distinct and independent mechanosensory accumulation of β_H_-Spec and MyoII. In the control cell (top row), MyoII was enriched at the tip of the aspirated portion of the cell (arrowhead, top row, middle panel) and β_H_-Spec enriched at the base (arrowheads, top row right panel). Suppressing MyoII activity by Y27632 abolished the accumulation of MyoII (middle row, middle panel), but not β_H_-Spec (arrowheads, middle row, right panel), whereas β_H_-Spec knockdown (KD) did not affect MyoII accumulation (bottom row). Scale bars, 5 μm. (d) Quantification of protein accumulation at the base of aspirated cells. Background-subtracted fluorescent intensities at the base (I_b_) and at the opposite pole of the cell body (I_o_) were measured, and the ratio (I_b_/I_o_) was calculated. (e) Quantification of MyoII accumulation at the tip of aspirated cells. Background-subtracted fluorescent intensities at the tip (I_t_) and at the opposite pole of the cell body (I_o_) were measured, and the ratio (I_t_/I_o_) was calculated. In (d) and (e), each data point represents a cell and each horizontal line indicates the average ratio. (f-i) Coarse-grained simulation of deformation in the receiving cell induced by invasive protrusions. (f, left panel) Schematic graph of the geometry of a protrusion on the surface of a receiving cell. (f and g) Heat maps of the simulated dilation (f, middle panel and g, left panel) and shear (f, and g, right panels) deformations induced by protrusions with a radius (*r_0_*) of 2.5 μm (F’ and F”) and 200 nm (G and G’), respectively. (h and i) Plots of dilation and shear deformation along protrusions with an *r_0_* of 500 nm (H) and 200 nm (I). Each data point is the average deformation of 4 different lines along the protrusions. Note that the shear deformation at the tip region increases in (i) compared to (h). (j) Dilation and shear deformations at the tip area (0.0-0.2) of protrusions with different *r_0_*. Note the gradual increase in shear deformation at the tip with a decrease in the protrusion radius. The length (l) of all protrusions is 2 μm. (k and l) β_H_-Spec accumulation in response to mechanical stress revealed by AFM. (k) Schematic drawing of the AFM experiments. A cantilever applied a pushing force to the cortex of an S2R+ cell. (l) Stills from live imaging of an S2R+ cell expressing GFP-β_H_-Spec and RFPMyoII. Both β_H_-Spec and MyoII rapidly accumulated to the indented area generated by the cantilever (arrowheads). Scale bar, 10 μm.

It is intriguing that α/β_H_-spectrin and MyoII show distinct domains of mechanosensitive accumulation revealed by MPA. Previous coarse-grained modeling suggested that the tip of an aspirated cell corresponds to an area of maximal actin network dilation (or radial expansion), whereas the base area is under maximal shear deformation (or shape change)^29^. These analyses suggest that MyoII is a mechanosensory protein for actin network dilation, whereas α/β_H_-spectrin responds specifically to shear deformation. Consistent with the distinct areas of mechanosensitive accumulation of MyoII and spectrin, β_H_-spectrin remained accumulated to the base area in cells treated with Y27632, a small molecule that decreases MyoII activity by inhibiting MyoII’s upstream activator ROCK (compare Fig. 3c, top and middle row; Fig. 3d), and MyoII remained accumulated at the tip of β_H_-spectrin knockdown cells (compare Fig. 3c, top and bottom row; Fig. 3e).

The distinct domains of mechanosensitive accumulation of MyoII and spectrin induced by pulling forces prompted us to ask whether they exhibit a similar response to pushing forces. Course-grained modeling of cells invaded by protrusions with a 5-μm diameter predicted clear separation of dilation vs. shear domains along the invasive protrusion, with maximal dilation at the tip area and maximal shear deformation at the base (Fig. 3f). However, when the invasive protrusions became thinner (approaching a diameter of 400 nm), there was a gradual increase of shear deformation at the tip area, where the dilation deformation remained largely the same (Fig. 3g-j). This model predicted that the mechanosensitive accumulations of spectrin and MyoII induced by thin protrusions may overlap at the tip area, although the maximal shear deformation still resided at the base. To test this directly, we performed atomic force microscopy (AFM) experiments, in which a pushing force was applied to cells expressing GFP-β_H_-spectrin and RFP-MyoII by a cantilever with a diameter of ~200 nm, closely mimicking the length scale of the invasion protrusions at the fusogenic synapse (Fig. 3k). β_H_-spectrin and MyoII exhibited rapid and largely overlapping domains of accumulation to the indented area by the cantilever (Fig. 3l; Supplementary Video 4), validating the pattern of mechanosensitive response predicted by the course-grained model. Consistent with this, both β_H_-spectrin and MyoII accumulated in largely overlapping domains together with Duf at the fusogenic synapse in founder cells of *Drosophila* embryos (Fig. 1k-m)^6^.

What is the biological function of spectrin accumulation at the fusogenic synapse? In *α/β_H_-spectrin* double mutant embryos, the founder cell-specific adhesion molecule Duf and its interacting protein Ants were both dispersed along the muscle cell contact zone between two fusion partners, instead of forming a tight cluster at the fusogenic synapse as in wild-type embryos (Fig. 4a, b; Extended Data Fig. 5a, b). Time-lapse imaging revealed dynamically dispersing Duf at the fusogenic synapse in the *α/β_H_-spectrin* mutant embryos (Supplementary Video 5), compared to the tight Duf cluster associated with dense F-actin foci in wild-type embryos (Fig. 4c; Supplementary Video 6). Occasionally, aggregated Duf clusters were initially observed in these mutant embryos, but in these cases Duf gradually diffused over time, suggesting that spectrin is required for the maintenance, but not the initiation, of the Duf cluster (Fig. 4d; Supplementary Video 5).

**Figure 4.**
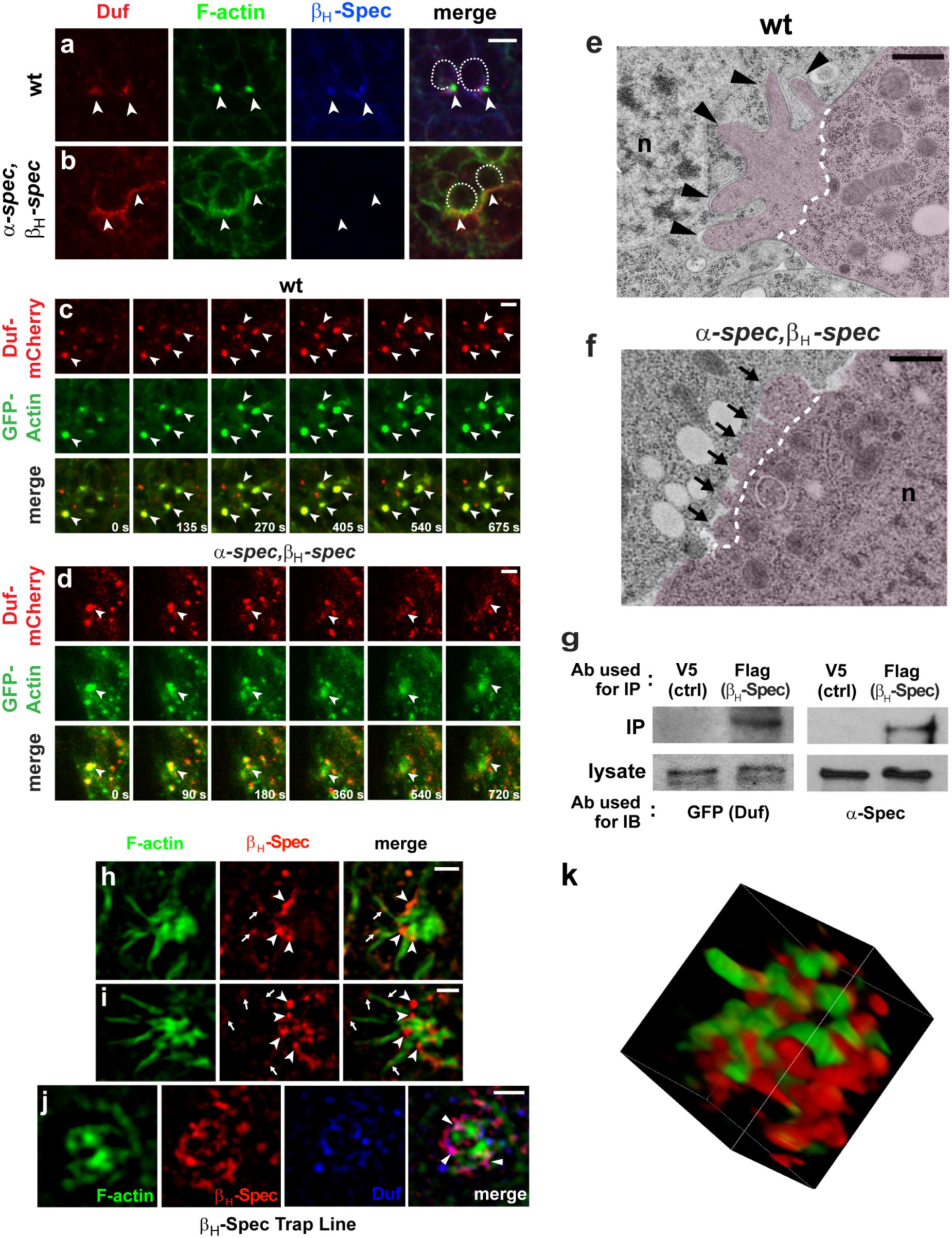
The spectrin network restricts cell adhesion molecules and constricts invasive protrusions at the fusogenic synapse. (a-b) Duf is dispersed at the fusogenic synapse in *α/β_H_-spectrin* double mutant embryos. Stage 14 embryos labeled with anti-Duf (red), phalloidin (F-actin) and anti-β_H_-Spec (blue). (a) Duf is concentrated in a tight cluster at the fusogenic synapse (arrowheads) with the dense F-actin focus in a wt embryo. (b) Duf is dispersed along the muscle cell contact zone (arrowheads) with fuzzy actin filaments in a *α/β_H_-spectrin* mutant embryo. FCMs are outlined in the merge panels. Scale bar, 5 μm. (c-d) Stills from time-lapse movies of wt (c) or *α/β_H_-spectrin* mutant (d) embryos expressing Duf-mCherry (red) and GFP-actin (green). Arrowheads indicate fusogenic synapses. Note that Duf remained in a tight cluster associated with the dense F-actin foci in wt, but diffused over time accompanied by the dispersal of the F-actin focus in an *α/β_H_-spectrin* mutant embryo. Scale bar, 5 μm. (e-f) Electron micrographs of the invasive PLS in wt (e) and *α/β_H_-spectrin* mutant (f) embryos. FCMs were pseudo-colored in purple. Dash lines delineate the F-actin enriched area of the PLS. Note the long, thin and well separated fingerlike protrusions in wt (arrowheads in e) and the stubby and closely abutting toe-like protrusions in *α/β_H_-spectrin* mutant (arrows in f) embryos. n, nucleus. Scale bars, 500 nm. (g) Biochemical interaction between β_H_-Spec and Duf. Co-immunoprecipitation (IP) experiment using extracts from embryos expressing Flag-β_H_-Spec and Duf-GFP. Duf was pulled down with anti-Flag antibody, suggesting that β_H_-Spec interacts with Duf. α-Spec was detected to indicate the presence of β_H_-Spec, the latter of which was difficult to detect due to its large size (~480 kDa). (h-j) The spectrin network constricts the actin-propelled protrusions. Structural illuminated microscopy (SIM) images of fusogenic synapses, labeled with F-actin (green) and β_H_-Spec (red). (h and i) Side-by-side view of two distinct fusogenic synapses between an FCM (right) and a founder cell (left). Note that the long and thin actin-propelled protrusions from the FCM penetrated through spectrin-free domains, but not spectrin-enriched patches (arrowheads), within the network. The long and thin protrusions triggered additional accumulation of spectrin at the tips or along the protrusions (arrows). (j) Top-to-bottom view of a fusogenic synapse. Note that the F-actin enriched areas were non-overlapping with those of β_H_-Spec or Duf (blue). Also note the significant overlapping domains of β_H_-Spec and Duf (arrowheads). (k) 3D reconstruction of the fusogenic synapse, highlighting the mutually exclusive localization of F-actin and β_H_-Spec.

Since the cell adhesion molecules from the two fusion partners, Duf and Sns, interact in trans at the fusogenic synapse^30^, we tested whether Duf dispersal in founder cells of *α/β_H_-spectrin* mutant embryos affects Sns distribution in the FCM in a cell non-autonomous manner. Indeed, Sns was also dispersed along the muscle cell contact zone at the fusogenic synapse (Extended Data Fig. 5, compare c and d). Moreover, the actin nucleation promoting factors and their interacting proteins, such as the WASP-interacting protein (known as Solitary or Sltr), which are recruited by Sns to the fusogenic synapse^31^, were dispersed in these mutant embryos (Extended Data Fig. 5, compare e and f). Consequently, the F-actin structure generated by the FCM appeared diffused (Fig. 4b; Extended Data Fig. 5b, d, f), with an average fluorescent intensity of 61 ± 19 per focus on a 0-255 scale (n=35), compared to 170 ± 15 per focus (n=28) in wild-type embryos. Electron microscopy studies revealed stubby and closely abutting toe-like protrusions in α/β*_H_-spectrin* mutant, compared to the long, thin and well separated fingerlike protrusions in wild-type embryos (Fig. 4e, f)^4,17,18^. Thus, Duf restriction by α/β_H_-spectrin in founder cells non-autonomously regulates Sns localization and the distribution of polymerized actin filaments along the cell contact zone at the fusogenic synapse in FCMs.

To investigate how spectrin maintains the Duf concentration at the fusogenic synapse, we performed co-immunoprecipitation experiments using *Drosophila* embryos expressing Flag-β_H_-spectrin and Duf-GFP specifically in the mesoderm. Antibody against Flag, but not a control antibody, co-precipitated both Duf-GFP and α-spectrin, suggesting that α/β_H_-spectrin restricts Duf via biochemical interactions (Fig. 4g). In the absence of spectrin-Duf interaction, such as in *duf,rst* mutant embryos expressing DufΔintra, even though the F-actin foci formed initially due to the trans-interaction between DufΔintra and Sns, the condensed foci eventually dispersed at the fusogenic synapse (Supplementary Video 7), similar to the phenotype of the *α/β_H_-spectrin* mutant embryos (Fig. 4a, b; Extended Data Fig. 5b, d, f; Supplementary Video 5). Thus, the intracellular domain of Duf is required for stabilizing the mechanosensitive accumulation of spectrin, likely through spectrin-Duf interactions.

Besides restricting cell adhesion molecules, the abnormal morphology of the invasive protrusions in *α/β_H_-spectrin* mutant embryos suggests that spectrin may play an additional role in shaping these intercellular protrusions. Using structured illumination (SIM) microscopy, we observed that the spectrin accumulation triggered by initial invasive protrusions did not form an evenly spaced network like in red blood cells and neuronal axons^8-10^. Instead, the network contained spectrin-enriched patches interspersed with spectrin-free domains (Fig. 4h-k). Interestingly, actin-propelled membrane protrusions only penetrated through the spectrin-free domains but not spectrin-enriched patches (Fig. 4h-k; Supplementary Video 8). The long and thin protrusions triggered additional mechanosensitive accumulation of spectrin at the tip and along the side (Fig. 4h, i), which may constrict further protrusions. These results suggest that the spectrin network in the founder cell functions as a “cellular sieve” to restrict the diameters of the invasive protrusions from the FCM. The resulting long and thin protrusions put the fusogenic synapse under high local tension and promote plasma membrane fusion^4,6^.

Taken together, this study has revealed a dynamic mechanosensitive response of spectrin and its cell non-autonomous function in shaping the architecture of invasive protrusions. During cell-cell fusion, the mechanosensitive accumulation of spectrin in the receiving fusion partner sets up a transient spectrin-enriched network at the fusogenic synapse, which, in turn, restricts cell adhesion molecules via biochemical interactions and constricts the invasive protrusions to increase local tension and drive the fusion process (Extended Data Fig. 6). What triggers the mechanosensitive accumulation of spectrin? Previous studies using *in vitro* reconstituted actin networks have demonstrated that mechanical loading/compression increases the density of the actin network^32^. We suggest that the invasive protrusions at the fusogenic synapse compress and deform the actin network in the receiving fusion partner, thus generating numerous additional spectrin binding sites and leading to its mechanosensitive accumulation. The α/β_H_-spectrin bound to the dynamic actin network, in turn, may transiently stabilize the network by unfolding the spectrin repeats, as demonstrated in red blood cells under shear stress ^33^. What makes spectrin dynamic at the fusogenic synapse? In many previously described cellular contexts where spectrin forms a static membrane skeletal structure, the spectrin-actin interaction is stabilized by accessory proteins, such as adducin^34^ and protein 4.1^1,2,35,36^. Interestingly, adducin and protein 4.1 were both absent at the fusogenic synapse (Extended Data Fig. 1b, c), which may account for the dynamic behavior of α/β_H_-spectrin during the fusion process. It is worth noting that the fusion-promoting function of spectrin is not fly-specific; it is conserved in mouse myoblast fusion (Extended Data Fig. 7). Given the widespread expression of spectrin in most eukaryotic cell types, our characterization of spectrin as a dynamic mechanosensory protein in fusogenic cells has broader implications for understanding spectrin functions in many dynamic cellular processes beyond cell-cell fusion.

## ACKNOWLEDGEMENTS

We thank the Bloomington stock center for fly stocks, B. Paterson for the MHC antibody, Guofeng Zhang for help with HPF/FS, J. Nathans for sharing confocal microscopes, and D. Pan for critically reading the manuscript. This work was supported by NIH grants (R01 AR053173 and R01 GM098816), American Heart Association Established Investigator Award and HHMI Faculty Scholar Award to E.H.C.; NIH grants (R01 GM66817 and R01 GM109863) to D.N.R; NIH grants (R01 GM074751 and R01 GM114671) to D.A.F; NSF grant #MCB-1122013 to G.T.; and NSFC grant #11572316 to T.L. R.D. was supported by an AHA postdoctoral fellowship; K.S. by an AHA Scientist Development Grant; S.S. by a Life Sciences Research Foundation postdoctoral fellowship; and D. L. by a Canadian Institute of Health Research postdoctoral fellowship.

## AUTHOR CONTRIBUTIONS

R.D. initiated the project. R.D., J.K., K.S. and E.C. planned the project, performed experiments in Figures 1, 2, 4 (a-d, g) and Extended Data Figures 2, 3, 5, 7, and discussed the data. J.K. and E.C. collaborated with E.S. and D.R. on the MPA experiments and with S.S. and D.F. on the AFM experiments. D.L carried out the SIM experiments. S.L. carried out the EM experiments. T.L. contributed the coarse-grained models. G.T. contributed a spectrin construct. R.D., J.K., K.S. and E.C. made the figures. J.K and E.C. wrote the paper. All authors commented on the manuscript.

## AUTHOR INFORMATION

The authors declare no competing financial interests.

## METHODS

### Fly genetics

Fly stocks used in this study: *α-spec^rg41^* (*α-spec* mutant; Bloomington Stock Center), *kst^14.1^* (*β_H_-spec* mutant)^19^, *Df(3L)1226* (*β_H_-spec* deficiency line)^19^, *UAS-mini-β_H_-Spec*^37^, *kst^MI03134-GFSTF.1^*(*β_H_-spec* trap line; Bloomington Stock Center), *dpak3^zyg(del)^* ^18^, *sltr^S1946^* ^31^, *UAS-DufΔintra-Flag*^38^, *sns-GAL4*^39^, *rP298-GAL4*^24^, *UAS-GFP-Actin* and *UAS-Actin-mRFP* (Bloomington Stock Center). Transgenic flies carrying *UAS-V5-β_H_-Spec*, *UAS-mCherry-β_H_-Spec*, *UAS-Duf-GFP* and *UAS-Duf-mCherry* were generated by P-element-mediated germline transformation. To express genes in muscle cells, females carrying the transgene under the control of an *UAS* promoter were crossed with *twi-GAL4* (all muscle cells), *rP298-GAL4* (founder cells) and *sns-GAL4* (FCMs) males, respectively. *α/β_H_*-spectrin double mutant, *α-spec^rg41^,kst^14.1^/TM6* (labeled as *α-spec,β_H_-spec* in figures), was generated using standard genetic methods.

### Molecular biology

Full-length *β_H_-spec* was amplified by PCR (with or without a tag) from cDNAs generated from mRNA of stage 11-15 *w^1118^* flies. Due to the large size of *the β_H_-spec* gene, three fragments were individually amplified using the primers as follows:

1. β_H_-spec-5’: GACCGGTCAACATGACCCAGCGGGACGGCATC
2. β_H_-spec-3721-3’: CTCCACGAATTCGGTGTCATG
3. β_H_-spec-3721-5’: CATGACACCGAATTCGTGGAG
4. β_H_-spec-8214-3’: CTCACCCTCTAGAATGCTATTG
5. β_H_-spec-8214-5’: CAATAGCATTCTAGAGGGTGAG
6. β_H_-spec-3’: CCAAGCGGCCGCTCACTGTGGCGGGACTTGACTC

The three PCR fragments were then subcloned into a *Drosophila* transformation vector pUAST. To generate the UAS-*β_H_-Spec*ΔN and UAS-*β_H_-Spec*ΔC constructs, the following primers were used:

1. β_H_-spec-3865-5’: GGAATTCCAACATGGTGTGTCGATCTGCAAATGTTC
2. β_H_-spec-8028-3’: GGTCTAGATCACAGCTGATGGGCCTCAGTTAG

To generate pDEST-*β_H_-Spec* constructs for GST-fusion proteins for the F-actin co-sedimentation assays, the Gateway cloning system was used (Invitrogen)with the following primers:

1. β_H_-spec-1-C: GGGGACAAGTTTGTACAAAAAAGCAGGCTTCATGACCCAGCGGGACGGCATC
2. β_H_-spec-1-K: GGGGACCACTTTGTACAAGAAAGCTGGGTTTTACTTCTTGCGATCTGCGTCCAT
3. β_H_-spec-29-C: GGGGACAAGTTTGTACAAAAAAGCAGGCTTCGGAGCCAAACAAGTCCACGTC
4. β_H_-spec-31-K: GGGGACCACTTTGTACAAGAAAGCTGGGTTTTATTGGGACGCCGCATTCTGGCG
5. β_H_-spec-34-C: GGGGACAAGTTTGTACAAAAAAGCAGGCTTCCCGAACATGCAACTGCTTAGC
6. β_H_-spec-34-K: GGGGACCAC TTTGTACAAGAAAGCTGGGTTTTATCACTGTGGCGGGACTTGACT dsRNAs were synthesized by *in vitro* transcription with gene-specific primers containing the T7 promoter sequence (TTAATACGACTCACTATAGGGAGA) at the 5’ end (MEGAscript; Ambion). Synthesized dsRNAs were purified using NucAway™ Spin Columns (Ambion).

### Immunofluorescent staining and imaging

Fly embryos were fixed and stained as described previously^4,31^. The following primary antibodies were used: rabbit α-muscle myosin heavy chain (1:1000) (gift from B. Paterson), rabbit anti-β_H_-Spectrin (1:100)^19^, rabbit anti-β-Spectrin (1:400)^40^, mouse anti-α-Spectrin (1:1; DSHB), guinea pig α-Duf (1:500)^4^, guinea pig α-Ants (1:1000)^23^, rat α-Sltr (1:30)^31^, rat α-Sns (1:500)^41^, mouse anti-adducin (1:400; DSHB), mouse anti-protein 4.1 (1:400; DSHB), mouse α-Flag (1:200; Sigma) and mouse α-V5 (1:200; Invitrogen). The following secondary antibodies were used at 1:200: Alexa488- (Invitrogen), Cy3-, and Cy5- (Jackson Laboratories) conjugated and biotinylated (Vector Laboratories) antibodies made in goats. For phalloidin staining, FITC- or Alexa568-conjugated phalloidin (Invitrogen) were used at 1:200. Fluorescent images were obtained on an LSM 700 Meta confocal microscope (Zeiss), acquired with LSM Image Browser software (Zeiss) and Zen software (Zeiss), and processed using Adobe Photoshop CS. For quantification of fluorescent signals, the signal intensities of cellular area of interest and control area were measured and processed for presentation by the Image J program (<u>http://imagej.nih.gov/ij/</u>).

### *Drosophila* cell culture

S2 and S2R+ cells were cultured in Schneider’s medium (Gibco) supplemented with 10% fetal bovine serum (FBS) (Gibco) and penicillin/streptomycin (Sigma). Cells were transfected using Effectene (Qiagen) per the manufacturer’s instructions. For RNAi knockdown, cells were first incubated with 3-5 μg/ml of dsRNA for 2-4 days and then transfected with appropriate DNA constructs.

For immunofluorescent staining, cells were fixed with 4% formaldehyde in PBS, washed in PBST (PBS with 0.1% Triton X-100) and PBSBT (PBST with 0.2% BSA) consecutively, and stained with the following antibodies in PBSBT: mouse α-V5 (1:2000; Invitrogen) and rabbit α-GFP (1:1000; Invitrogen). Secondary FITC-, Cy3-, or Cy5-conjugated antibodies were used at 1:400 (Jackson Immunoresearch). To visualize F-actin, FITC- or Alexa 568-conjugated phalloidin (Invitrogen) was used at 1:500 in PBST.

### Mouse C2C12 myoblast culture

A pair of predesigned siRNAs against the mouse Sptbn5 gene and negative control siRNA were obtained from Qiagen. RNAi was performed per manufacturer’s instructions. Briefly, approximately 3x10^5^ cells were seeded on each well of a 6-well tissue culture dish and incubated overnight. On day 2, cells were transfected with the mix of two siRNAs against Sptbn5 (200 pmol each) using HiPerFect transfection reagent (Qiagen). On day 3, the cells were washed and differentiated, and cells that were treated in parallel were subjected to RT-PCR to access knock-down (KD) level. Fusion index was determined when the KD level was more than 80% as follows. Three days post-differentiation, cells were fixed and stained with anti-MHC antibodies (MF20) to identify differentiated cells and with CellMaskTM Orange membrane stain (Molecular Probes, Invitrogen) to visualize the overall cell/syncytia morphology. Cells were mounted using Prolong Gold antifade reagent with DAPI (Molecular Probes, Invitrogen) to visualize the nuclei. Ten random 40X microscopic fields under Axioscope 2 (Zeiss) were counted for each experiment and repeated in at least three independent experiments (total number of fields ≥30). The fusion index was calculated as a mean percentage of nuclei in syncytia versus total number of nuclei per field in differentiated cells.

### Time lapse imaging and fluorescent recovery after photobleaching (FRAP)

Time lapse imaging of embryos was performed as previously described^4^. Briefly, embryos expressing fluorescently tagged proteins in muscle cells were collected and dechorionated in 50% bleach. Subsequently, embryos were washed in water, placed onto a double-sided tape (3M), and covered with a layer of Halocarbon oil 700/27 (2:1; Sigma). Time lapse image acquisition was carried out on an LSM 700 Meta confocal microscope (Zeiss).

The FRAP experiments were performed as previously described^17^. A region of interest (ROI) was manually selected and imaged in 3 to 5 frames to record the original fluorescent intensity (pre-bleach). Then the ROI was quickly bleached to a level at lower than 20% of its original fluorescent intensity and subsequently imaged with an appropriate time interval (post-bleach). The fluorescent intensities of the pre- and post-bleached ROI were measured using the image J program. The Prism software was used to determine the maximum recovery level (the percentage recovery compared to the pre-bleach level) and the half-time of recovery (T_half_) using a kinetic curve fit with an exponential decay equation.

### Structured illumination microscopy (SIM)

Stage 13-14 embryos were fixed and stained as described above. The samples were then mounted in Prolong Gold (Molecular Probes) and imaged with an inverted microscope (Ti-E; Nikon) equipped with a 100x oil NA1.49 CFI SR Apochromat TIRF objective lens and an ORCAFlash 4.0 sCMOS camera (Hamamatsu Photonics K.K.). The images were acquired and processed with NIS-Elements AR software (Nikon).

### Electron microscopy

Embryos were fixed by the high-pressure freezing and freeze substitution (HPF/FS) method as previously described^4,42^. Briefly, a Bal-Tec device was used to freeze stage 12-14 embryos. Freeze-substitution was performed with 1% osmium tetroxide, 0.1% uranyl acetate in 98% acetone and 2% methanol on dry ice. Fixed embryos were embedded in Epon (Sigma-Aldrich) and cut into thin sections with an ultramicrotome (Ultracut R; Leica). The sections were mounted on copper grids and post-stained with 2% uranyl acetate for 10 min and Sato’s lead solution^43^ for 1 min to improve image contrast. Images were acquired on a transmission electron microscope (CM120; Philips).

### Recombinant protein purification and F-actin co-sedimentation assay

To purify GST-fused β_H_-Spec fragments from BL21-DE3 cells (NEB), protein expression was induced with 0.2 mM IPTG at room temperature for 12-15 hours. Cells were harvested and lysed by sonication in the lysis buffer: PBS (pH 7.4), 1% Triton X-100, 5 mM DTT, 1 mM PMSF, and complete-mini protease inhibitor cocktail (Roche). After centrifugation, the supernatant was collected and incubated with pre-equilibrated glutathione-agarose resin at 4°C for 2-3 hours. After washing in the lysis buffer, β_H_-Spec proteins were eluted with the Elution buffer: 50 mM Tris (pH 7.5), 150 mM NaCl, 5 mM DTT, 10 mM glutathione (Sigma).

F-actin co-sedimentation assay was performed following the manufacturer’s protocol (Cytoskeleton). Briefly, 0.5-1 μM purified proteins were incubated with 4 μM F-actin assembled from monomeric actin for 1 hour in F buffer: 5 mM Tris-HCl (pH 8.0), 0.2 mM CaCl_2_, 50 mM KCl, 2 mM MgCl_2_, 1 mM ATP. The F-actin/protein mixtures were centrifugated at 140,000 g for 30 min, and supernatants and pellets were separated and analyzed by SDS-PAGE and Coomassie Blue staining.

### Co-immunoprecipitation (Co-IP)

Embryos expressing GFP-tagged Duf and Flag-tagged β_H_-Spec (*kst^MI03134-GFSTF.1^/twi-GAL4; UAS-Duf-GFP/+*) were collected and dechorionated in 50% bleach. After being frozen in liquid nitrogen, embryos were disrupted in cold extraction buffer (50 mM Tris-HCl (pH 8.5), 150 mM NaCl, 0.5% sodium deoxycholate) with 20 strokes in a Dounce homogenizer. After centrifugation at 16,000 g for 20 min, supernatants were removed and incubated with either anti-V5 (control) or anti-Flag (β_H_-Spec) antibodies at 4°C for 2-3 hours. Protein-G-Sepharose beads (Roche) were used to precipitate antibodies. Precipitated proteins were analyzed by SDS-PAGE and western blotting.

### Micropipette aspiration (MPA)

Micropipette aspiration was performed as previously described^44^. Briefly, a pressure difference was generated by adjusting the height of a motor-driven water manometer. A fixed pressure of 0.4 nN/μm^2^ was applied instantly to the cell cortex of S2 cells with a polished glass pipette approximately 2-2.5 μm in radius. Images were collected in Schneider’s medium supplemented with 10% FBS on an Olympus IX81 microscope with a 40x (1.3 NA) objective and a 1.6x optivar, utilizing MetaMorph software and analyzed using ImageJ. After background correction, the fluorescence intensity at the sites of protein accumulation was normalized against the opposite cortex of the cell (I_b_/I_o_ or I_t_/I_o_, where I_b_, I_t_, and I_o_ is the intensity in the cortex at the base, tip, or opposite side of the cell being aspirated). An ANOVA with Fisher’s least significant difference was applied to determine statistical significance.

### Atomic force microscopy (AFM)

S2R+ cells were plated on glass coverslips coated with concanavalin A (Sigma, St Louis, MO) and transfected to express fluorescently tagged MyoII and β_H_-Spec using Effectene (Qiagen), per the manufacturer’s instructions. Lateral indentation experiments were conducted 2 days after transfection with a modified Catalyst AFM integrated with an Axio Observer fluorescence microscope (Zeiss). To determine the effect of a localized mechanical force on MyoII and β_H_-Spec localization, the cantilever (MLCT with a pyramidal tip, Bruker) was first brought into full contact, at around 50 nN setpoint force, with the glass surface on a cell-free area within 10 μm from a target cell. Next, the cell was laterally translated into the stationary cantilever using the piezoelectric XY stage and the NanoScope software (Bruker). The cantilever tip indented the edge of the cell by 2–5 μm. Cells were simultaneously imaged with a plan-apochromat 63x/1.4 NA oil immersion objective (Zeiss). Time lapse images were taken at 5 second intervals using the Micro-Manager software (http://micro-manager.org/wiki/Micro-Manager).

### Coarse-grained molecular mechanics modeling

In the coarse-grained model, the membrane-cortex composite is represented by a triangulated network where the nodes denote the crosslinking positions and the triangles resemble the meshes in the actin network, which is a network structure composed of proteins such as actin, actin crosslinkers, and MyoII. The system energy of the composite at the coarse-grained molecular level is calculated by

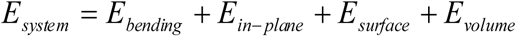

where *E_bending_* is bending energy from the plasma membrane; *E_in-plane_* is the in-plane elastic energy associated with the deformation of actin crosslinkers and the dilation of each mesh; *E_surface_* is the surface energy of the whole cell; *E_volume_* is the energy associated with the volume conservation of the cell^29,45,46^. The motion of the nodes is calculated by the Brownian dynamics equation (or overdamped Langevin dynamics equation) where the driving force is the derivative of the system energy with respect to the local position.

The initial configuration of the system was thermally annealed at room temperature until the fluctuation of the system energy was negligible. This configuration was then mapped to the shape of a receiving cell. For simplicity, the protrusion has a cylindrical shape with the radius *r_0_* and length *l* on the surface of a cell with a diameter of 10 μm. The tip of the protrusion was a spherical cap of a radius of *r_0_*. The final state of the system was achieved after 20 sec of Brownian dynamics simulation with a time-step of 10^-5^ sec. The area dilation of each node was determined by averaging of the dilation of the triangles with which the node of interest was associated. The shear deformation of each node was calculated in a similar way. The contour of the deformation on the protrusion was plotted by the software Tecplot. The deformations along the length direction of the protrusion were obtained by the extraction tool of the software.

---

**Extended Data Figure 1.**
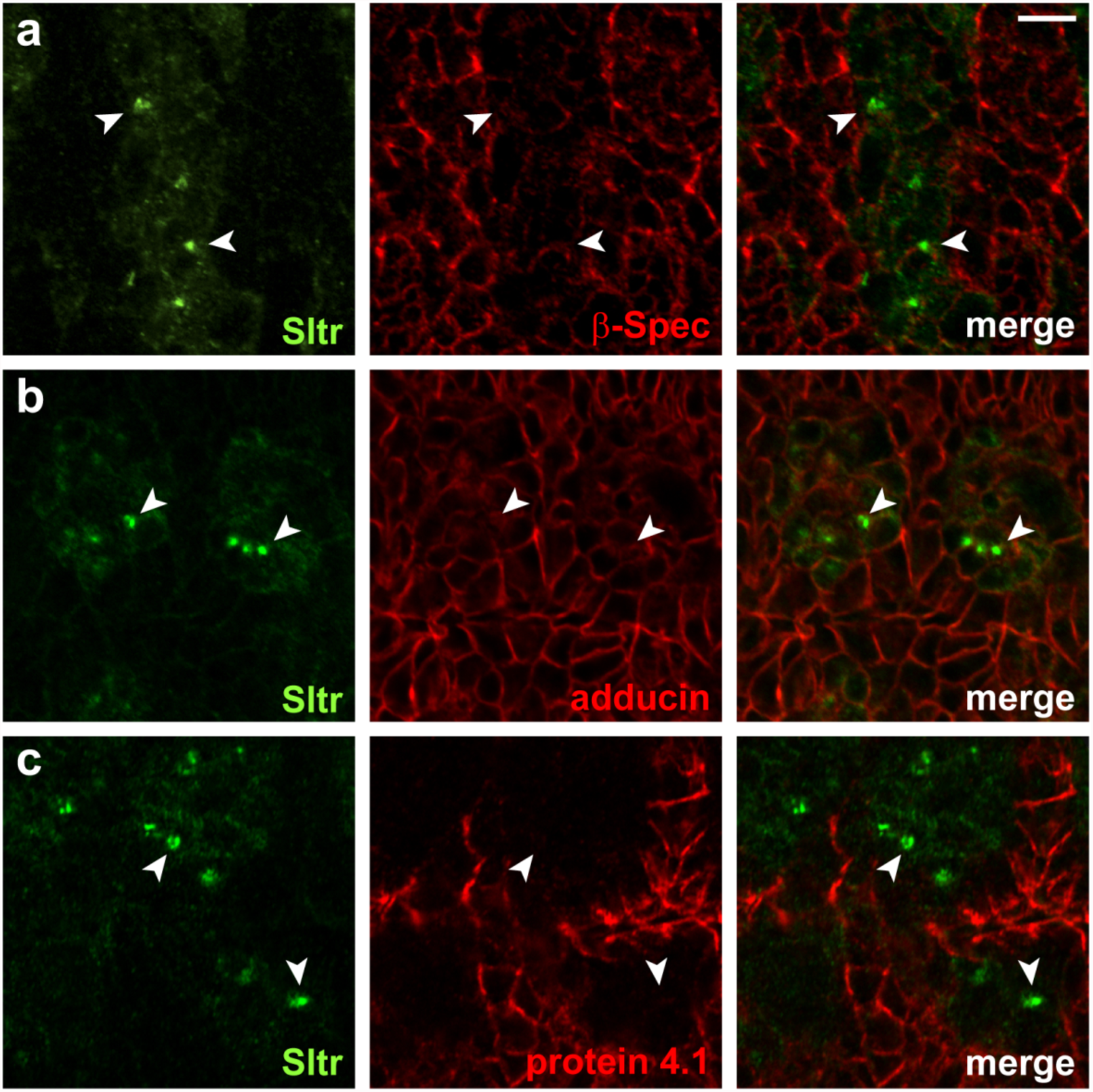
β-spectrin, adducin and protein 4.1 are not enriched at the fusogenic synapse. Fusogenic synapses in stage 13-14 wild-type embryos were marked by anti-Sltr staining (green; arrowheads). β-Spec (a), adducin (known as *hu-li tai shao* in *Drosophila*) (b), and protein 4.1 (known as *coracle* in *Drosophila*) (c) were visualized by immunostaining with respective antibodies. Note that none of these proteins was enriched at the fusogenic synapse. Scale bar, 5 μm.

**Extended Data Figure 2.**
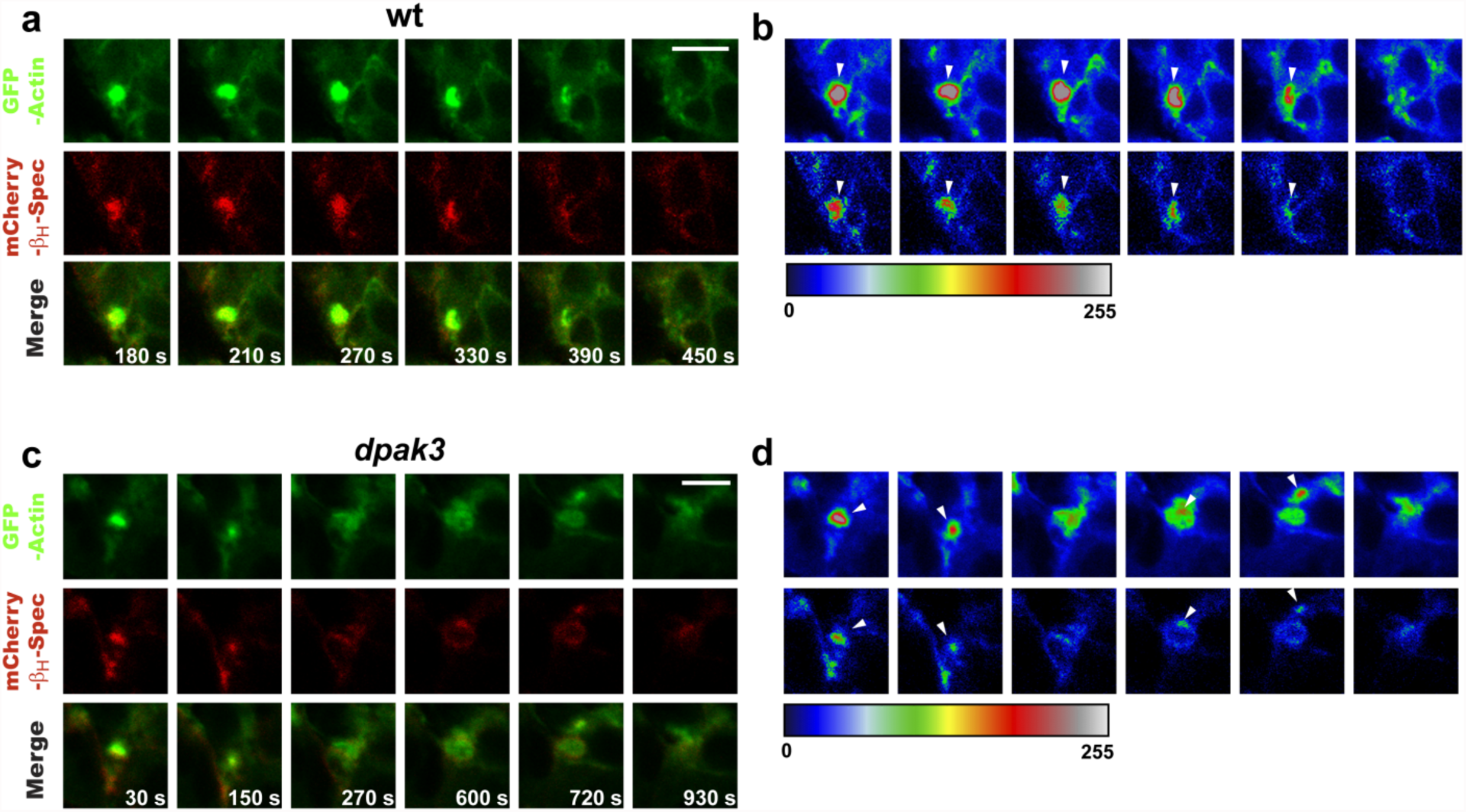
Dynamic accumulation of spectrin at the fusogenic synapse in response to PLS invasion. Stills from time lapse movies of wt (a) or *dpak3* mutant (c) embryos expressing GFP-actin (green) and mCherry-β_H_-Spec (red). The fluorescent intensities of F-actin foci and β_H_-Spec accumulation (a and c) were displayed by rainbow color-coded heat maps (b and d) on the arbitrary scale from 0 to 255. F-actin in *dpak3* mutant (c and d) was more diffused compared to the dense F-actin foci in wt (a and b). Note the dynamic changes in the intensity and morphology of β_H_-Spec accumulation correlating with the intensity of the F-actin foci (arrowheads in b and d). Scale bars, 5 μm.

**Extended Data Figure 3.**
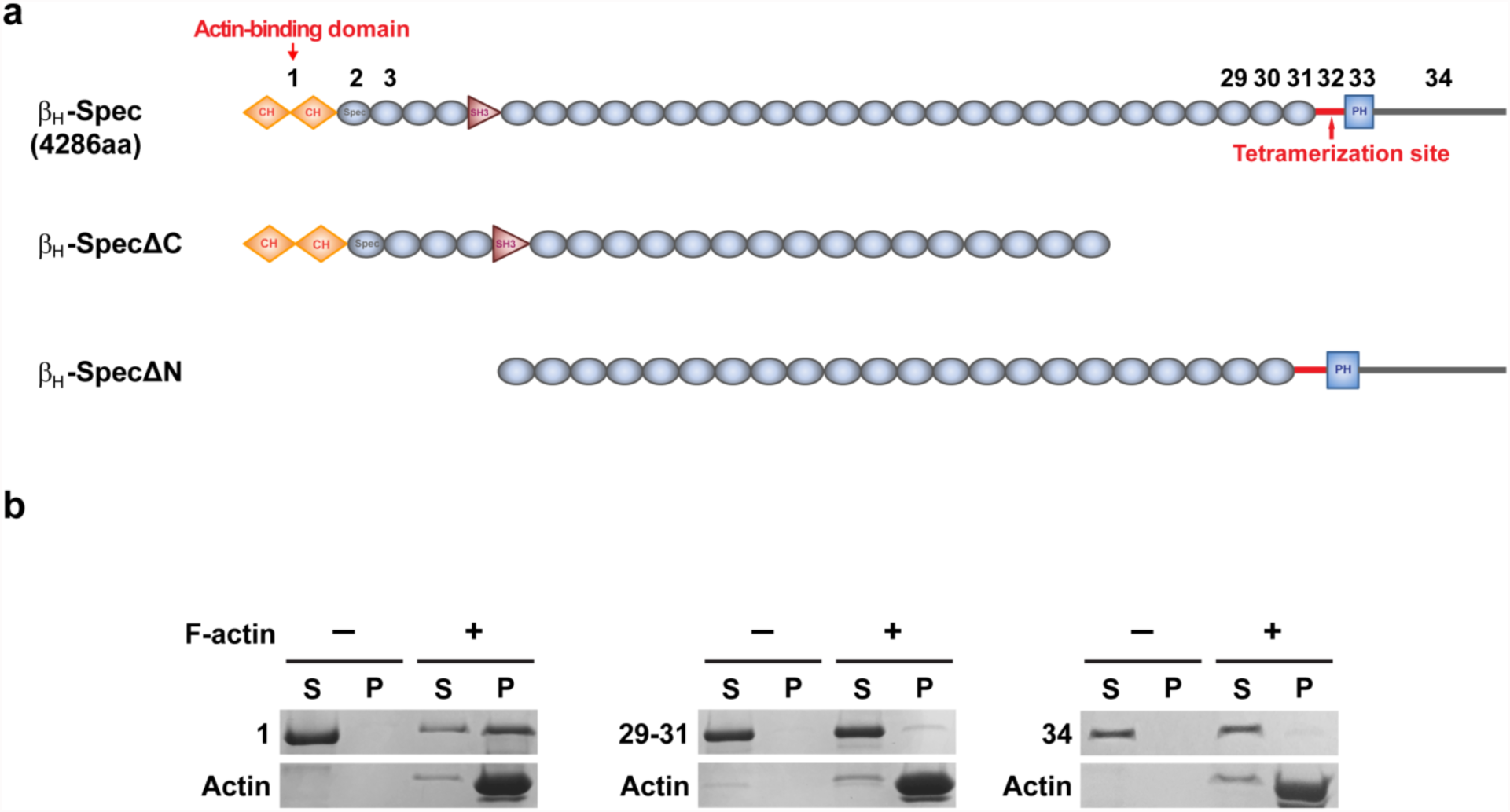
The N-terminal CH domains of βH-spectrin bind F-actin. (a) Domain structure of full length and mutant β_H_-Spec used in MPA and F-actin co-sedimentation assays. Actin-binding domains and the tetramerization site are indicated. Each distinct segment from the N-terminus to the C-terminus of β_H_-Spec is designated by a number. CH: calponin homology; SH3: Src homology 3; PH: pleckstrin homology; Spec: spectrin repeat. (b) F-actin co-sedimentation assay with purified β_H_-Spec fragments. The numbers (1, 29-31, 34) indicate β_H_-Spec fragments depicted in (a). S: supernatant; P: pellet. Note that β_H_-Spec-1 precipitated with F-actin, whereas β_H_-Spec-29-31 and β_H_-Spec-34 remained in the supernatant, confirming that the CH domains of β_H_-Spec bind F-actin.

**Extended Data Figure 4.**
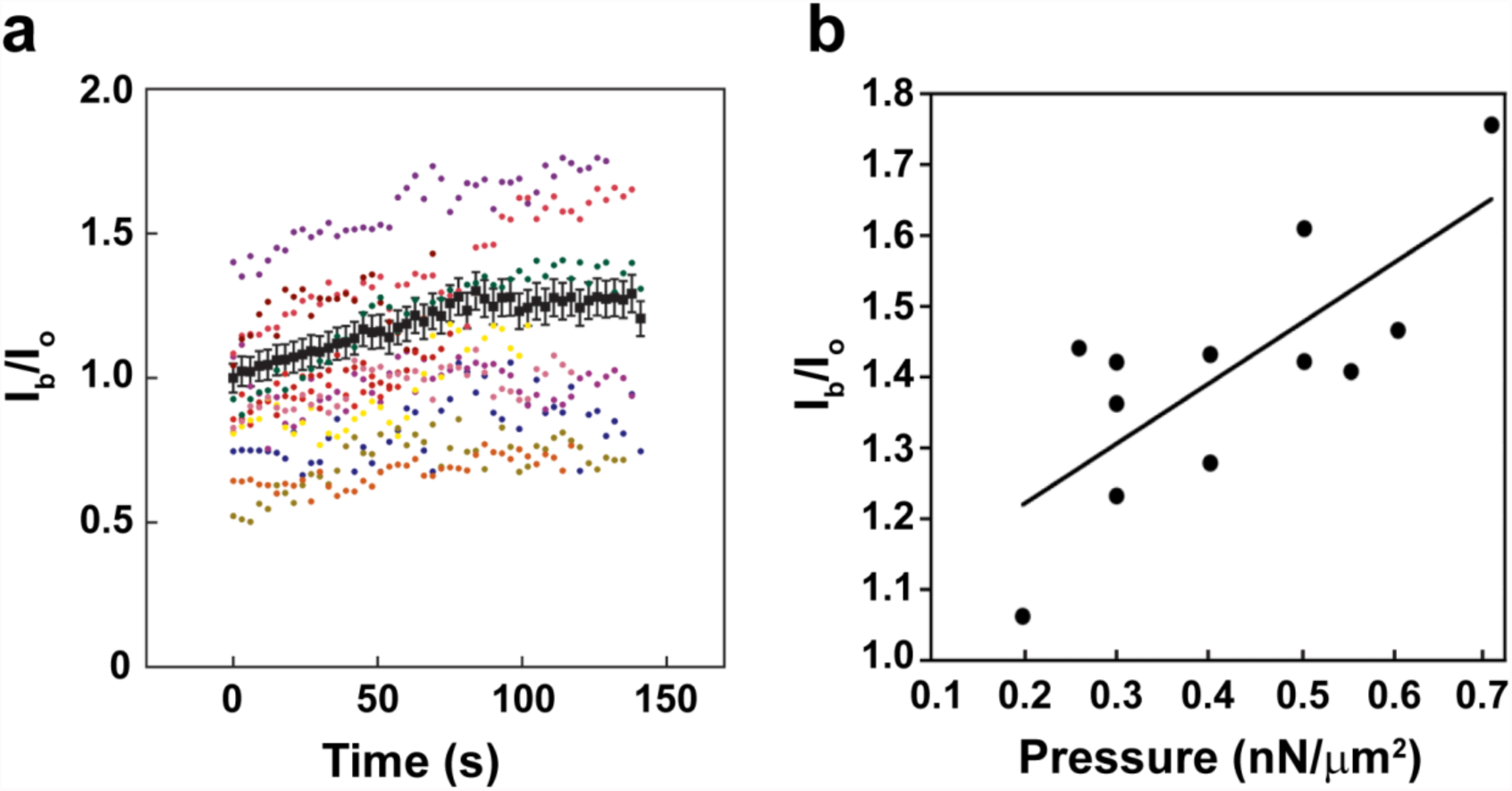
The mechanosensitive accumulation of spectrin is time- and force-dependent. (a) β_H_-Spec accumulation over time in MPA experiments. Different color codes indicate traces of individual cells examined. The average values at each time point are shown as black dots with error bars (SEM). β_H_-Spec accumulation linearly increased over time and reached its peak at around 80-90 sec after the onset of aspiration. (b) β_H_-Spec accumulation depends on applied force in MPA experiments. The increase of β_H_Spec accumulation is proportional to the elevation of applied pressure.

**Extended Data Figure 5.**
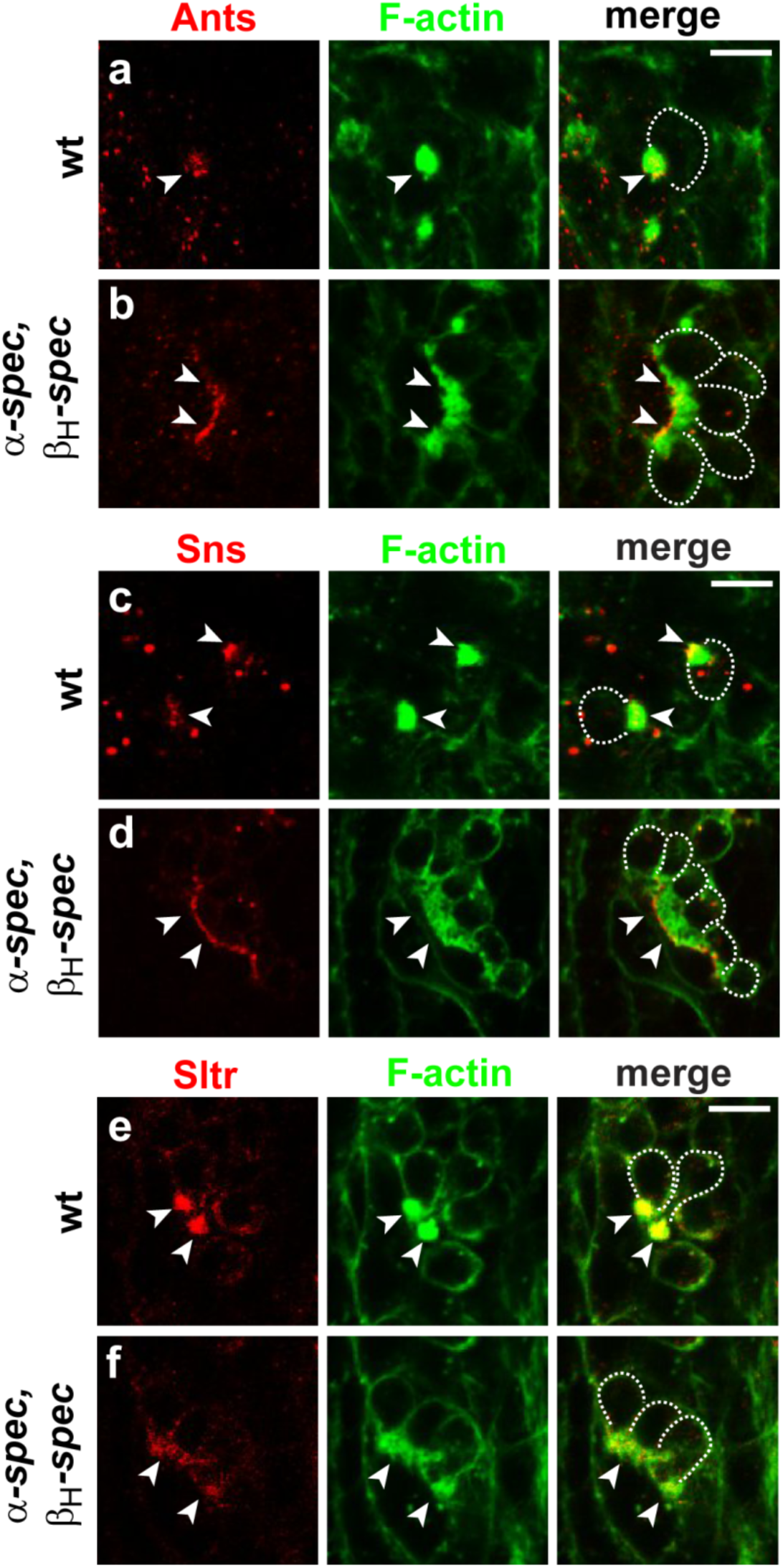
Ants, Sns, Sltr and F-actin are dispersed in *α/β_H_-spectrin* double mutant embryos. Fusogenic synapses (arrowheads) in wt and *α/β_H_-spectrin* mutant embryos were marked with F-actin foci (phalloidin staining). The localization of Ants (a-b), Sns (c-d), and Sltr (e-f) at the fusogenic synapse was visualized by immunostaining with respective antibodies. Note that all of these fusion-promoting proteins were dispersed in *α/β_H_-spectrin* mutant embryos (b, d, and f, bottom left panels) compared to their focal aggregation in wt embryos (a, c and e, top left panels). FCMs are outlined in the merge panels. Scale bars, 5 μm.

**Extended Data Figure 6.**
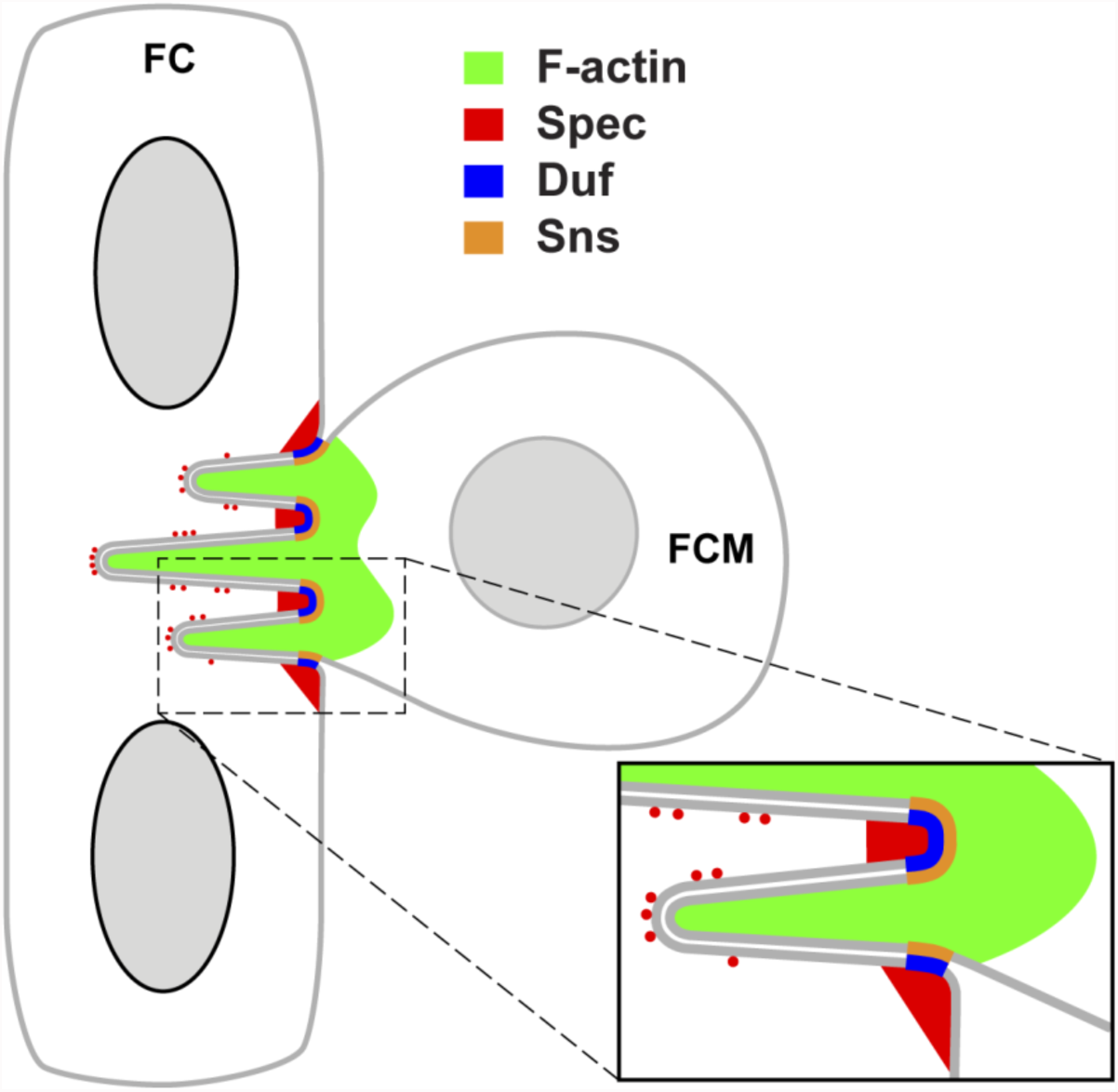
A working model of spectrin function at the fusogenic synapse. The spectrin network in the founder cell functions as a “cellular sieve” to constrict invasive protrusions from the FCM. The mechanosensitive accumulation of spectrin (Spec; red) at the fusogenic synapse is triggered by pushing forces generated by the initial invasive protrusions from the FCM. The accumulated spectrin (1) restricts the cell adhesion molecules, Duf (founder cell; blue) and Sns (FCM; gold), to the fusogenic synapse via biochemical interactions; and (2) forms an unevenly spaced network containing spectrin-enriched and spectrin-free microdomains. Actin-propelled membrane protrusions (F-actin; green) can only squeeze through the spectrin-free, but not spectrin-enriched, microdomains. The resulting fingerlike protrusions continue to trigger mechanosensitive accumulation of spectrin (red dots in the receiving cell), which may constrict new protrusions. These long and thin protrusions put the fusogenic synapse under high local tension and promote plasma membrane fusion.

**Extended Data Figure 7.**
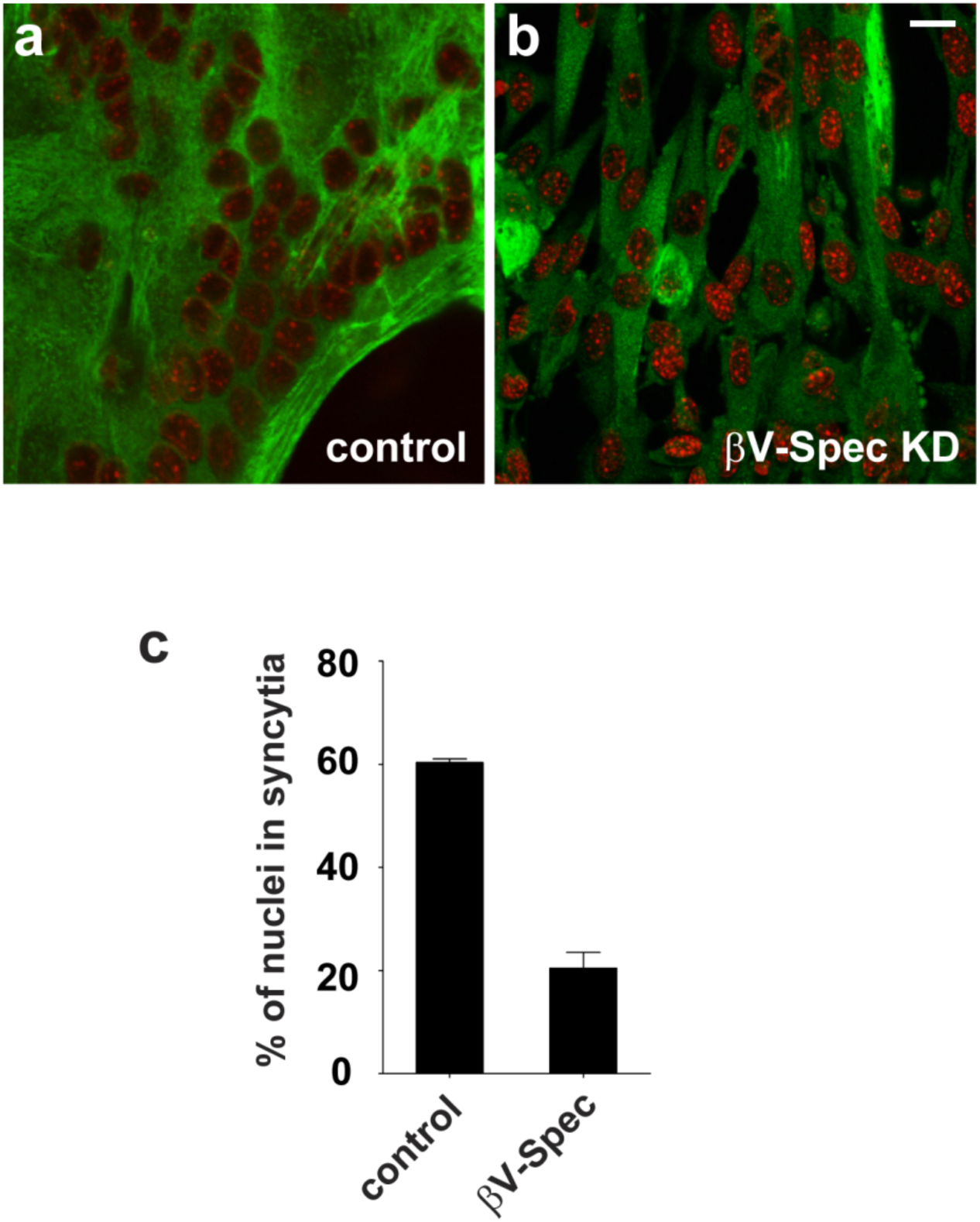
βV-spectrin, the mammalian homologue of *Drosophila* β_H_-spectrin, is required for C2C12 myoblast fusion. Confocal images of C2C12 cells treated with either control siRNA (a) or siRNA directed against Sptbn5 (βV-spectrin) (b). Cells were allowed to differentiate for 72 hours, then fixed and stained with anti-MHC (green) and DAPI (red) to visualize differentiated muscle cells. (c) The fusion index was calculated as the percentage of the number of nuclei in multinucleated syncytia versus the total nuclei in differentiated muscle cells (MHC-positive). In each experiment, cells were counted in 10 random 40X microscopic fields and the data were averaged from three independent experiments. Note that Sptbn5 knockdown (KD) significantly decreased the number of multinucleated syncytia. Error bar, standard deviation. Scale bar, 5 μm.

**Supplementary Video 1. Spectrin is dynamically associated with the F-actin focus at the fusogenic synapse**

Time-lapse imaging of muscle cells expressing GFP-actin (green) and mCherry-β_H_-spectrin (red) in a wild-type stage 14 embryo. The accumulation and dissolution of β_H_-Spec coincided with the appearance and disappearance of the F-actin foci.

**Supplementary Video 2. Spectrin accumulation correlates with the F-actin foci intensity**

Time-lapse imaging of muscle cells expressing GFP-actin (green) and mCherry-β_H_-spectrin (red) in a stage 14 *dpak3* mutant embryo. Arrow indicates a fusogenic synapse where the F-actin focus underwent dynamic aggregation and dispersal. Note that the intensity of β_H_-spectrin tightly correlated with that of the F-actin focus.

**Supplementary Video 3. Dynamic exchange of spectrin at the fusogenic synapse**

Fluorescent recovery after photobleaching (FRAP) of β_H_-spectrin at the fusogenic synapse (arrow). GFP-actin (green) and mCherry-β_H_-spectrin (red) were expressed in muscle cells in a stage 14 wild-type embryo. The mCherry fluorescence was photobleached and subsequently imaged every 30 sec. Note the rapid recovery of the mCherry signal after photobleaching.

**Supplementary Video 4. Mechanosensitive accumulation of spectrin and MyoII induced by lateral indentation using AFM**

Time-lapse imaging of an S2R+ cell co-expressing GFP-β_H_-spectrin (green) and RFP-Zipper (myosin heavy chain) (red). β_H_-spectrin and MyoII rapidly accumulated at the cell periphery laterally indented by a cantilever (arrow). Note the largely overlapping pattern of β_H_-spectrin and MyoII accumulation.

**Supplementary Video 5. Duf dispersal at the fusogenic synapse in** α**/**β***_H_-spectrin* double mutant embryos**

Time-lapse imaging of muscle cells expressing Duf-mCherry (red) and GFP-actin (green) in an α/β*_H_-spectrin* mutant embryo. Duf initially aggregated to a small cluster associated with a dense F-actin focus (arrowhead), but gradually dispersed together with F-actin over time.

**Supplementary Video 6. Duf resides in a tight cluster at the fusogenic synapse in wildtype embryos**

Time-lapse imaging of muscle cells expressing mCherry-Duf (red) and GFP-actin (green) in a stage 14 wild-type embryo. Note that Duf remained in a tight cluster associated with each dense F-actin focus (arrowheads).

**Supplementary Video 7. F-actin dispersal in the absence of the cytoplasmic domain of Duf**

Time-lapse imaging of muscle cells expressing mRFP-actin (red) and DufΔintra-Flag in a *duf,rst* mutant embryo. Note that F-actin initially formed a dense focus, but gradually dispersed over time. Arrowhead and arrow indicate two independent F-actin foci.

**Supplementary Video 8. 3D reconstruction of F-actin and spectrin at the fusogenic synapse based on SIM images**

Embryo was labeled with F-actin (green) and β_H_-Spec (red). Note the mutually exclusive localization of F-actin and β_H_-Spec.

